# MRI-related anxiety in healthy individuals, intrinsic BOLD oscillations at 0.1 Hz and heart rate variability in low frequency bands

**DOI:** 10.1101/359000

**Authors:** Gert Pfurtscheller, Andreas Schwerdtfeger, David Fink, Clemens Brunner, Christoph Stefan Aigner, Joana Brito, Alexandre Andrade

## Abstract

Participation in a MRI scan is associated with increased anxiety, thus possibly impacting baseline recording for functional MRI studies. We investigated in 23 healthy individuals without any former MRI experience (scanner-naïve) the relations between anxiety, 0.1-Hz BOLD oscillations and heart rate variability (HRV) in two separate resting state sessions (R1, R2). BOLD signals were recorded from precentral gyrus (PCG) and insula in both hemispheres. Phase-locking and time delays were computed in the frequency band 0.07–0.13 Hz. Positive (pTD) and negative time delays (nTD) were found. The pTD characterize descending neural BOLD oscillations spreading from PCG to insula and nTD characterize ascending vascular BOLD oscillations related to blood flow in the middle cerebral artery. HRV power in two low frequency bands 0.06–0.1 Hz and 0.1–0.14 Hz was computed. Based on the drop rate of the anxiety level from R1 to R2, two groups could be identified: one with a strong anxiety decline (large drop group) and one with a moderate decline or even anxiety increase (small drop group). A significant correlation was found only between the left-hemispheric time delay (pTD, nTD) of BOLD oscillations and anxiety drop, with a dominance of nTD in the large drop group. The analysis of within-scanner HRV revealed a pronounced increase of low frequency power between both resting states, dominant in the band 0.06–0.1 Hz in the large drop group and in the band 0.1–0.14 Hz in the small drop group. These results suggest different mechanisms related to anxiety processing in healthy individuals. One mechanism (large drop group) could embrace an increase of blood circulation in the territory of the left middle cerebral artery (vascular BOLD) and another (small drop group) translates to rhythmic central commands (neural BOLD) in the frequency band 0.1–0.14 Hz.

## Introduction

A major challenge associated with MRI scans performed with closed bore systems is the placement in a narrow place producing discomfort and even extensive fear. It is estimated that 25–37 % of patients experience moderate anxiety [1] and about 2 % intensive anxiety [2] during clinical scans. Although MRI-related anxiety and therewith activation of specific brain structures is common, little is known about their impact on slow intrinsic frequency fluctuations and resting state functional connectivity, respectively, in the frequency range near 0.1 Hz. A study on MRI-naïve young men with self-reported anxiety revealed highest anxiety levels during the first scan with a drop in the second scan [3]. MRI-related anxiety can affect the connectivity between the default mode network and left insula in youth and adults [4]. In a previous PLOSEONE paper about distinction between neural and vascular BOLD oscillations at 0.1 Hz [5] we reported on a significant hemispheric asymmetry during rest in healthy, scanner-naïve subjects, without giving a satisfied explanation about their origin. In the present paper we refer on additional results from this study on a cohort of healthy subjects with MRI-related anxiety, by taking into account the measured state anxiety and their change (drop rate) and offer an explanation for the dominant hemispheric asymmetry in the first resting state. Furthermore, relationships between heart rate variability (HRV) in two low frequency bands (0.06–0.1 Hz, 0.1–0.14 Hz) and anxiety drop rate are reported for the first time.

The discrimination between neural and vascular BOLD oscillations at 0.1 Hz is possible through computing the phase-locking value (PLV), either between two BOLD signals [5] or between BOLD and RRI signals [6,7]. By measuring phase-shifts of slow BOLD oscillations in precentral gyrus (PCG) and insula, and determining the sign of time delay (positive or negative) it is feasible to discriminate between ascending slow BOLD oscillations driven by the Mayer waves in cerebral blood flow and blood pressure (vascular BOLD) and descending BOLD oscillations most likely associated with neural activity fluctuations (neural BOLD; [5]).

There is an enduring interest in brain oscillations at 0.1 Hz and 10-s waves in blood pressure, heart rate and respiration, respectively, documented by two important papers published recently. One reports on entrainment of arteriole vasomotor and BOLD oscillations by gamma band activity [8] and the other on resonance breathing at a 10-s rate and “emotional well-being” [9]. The former contributes to the genesis of slow intrinsic neural BOLD oscillations near 0.1 Hz and the latter helps to explain the importance of low frequency HRV in emotion regulation [10, 11].

The goals of our paper are to report (i) on the discrimination between intrinsic neural and vascular BOLD oscillations near 0.1 Hz in anxious subjects based on only one pair of BOLD signals in each hemisphere, (ii) on the verification of two mechanisms related to of anxiety processing (cerebral blood circulation and central commands) and (iii) on the HRV in two low frequency bands (0.06–0.1 Hz, 0.1–0.14 Hz) during anxiety processing. The distinction between two bands close to 0.1 Hz may be of interest because power spectra of the heart rate intervals demonstrate very often two peaks centered at 0.8 Hz and 0.12 Hz [12].

## Methods

### Participants

From a total of 25 individuals (12 female) between 19–34 years (mean ± SD: 24 ± 3.2 years) two were excluded due to cardiac arrhythmia. All were naïve to the purpose of the study, had no former MRI experience, had normal or corrected-to-normal vision and were without any record of neurological or psychiatric disorders as assessed via self-report. All individuals gave informed written consent to the study protocol, which had been approved by the local Ethics Committee at the University of Graz.

### Experimental Protocol

The experimental task consisted of two resting states (R1, R2) and two within-scanner questionnaires on state anxiety (AS1, AS2) separated by about 30 minutes. The task started with the first questionnaire (AS1) and was followed by the first resting state (R1). Thereafter, two movement tasks (each lasting 600 s) were performed. The session ended with the second resting state (R2) and second questionnaire (AS2). Filling out each questionnaire took approximately 5 minutes and each resting state lasted for about 350 s. Individuals were requested to keep their eyes open, to stay awake, and to avoid movements during the resting states.

State anxiety was assessed with the state version of the state-trait anxiety and depression inventory (STADI; [13], which was presented on a screen within the scanner. The STADI is an instrument constructed to assess both state and trait aspects of anxiety and depression. It is based on the State-Trait Anxiety Inventory [14], but allows a reasonable separation of anxiety and depression symptoms. Items were answered with a trackball following each resting state.

### fMRI data acquisition and preprocessing

Functional images were acquired on a 3 T scanner (Magneton Skyra, Siemens). A multiband GE-EPI sequence [15] was applied with the following parameters: multiband factor 6, voxel size 2×2×2 mm^3^, TR/TE=871/34 ms, flip angle 52 degrees, matrix 90×104, 66 contiguous axial slices, FOV=180×208 mm^2^. Pre-processing and region of interest (ROI) signal extraction was performed using the DPARSF toolbox [16]. Pre-processing included the removal of the first 10 volumes, slice-timing correction adapted for multiband acquisitions, rigid-body motion correction, normalization to Montreal Neurological Institute (MNI) space, resampling to 2-mm isotropic voxels, spatial smoothing with a 4-mm FWHM Gaussian kernel and linear detrending. Lastly, the BOLD time courses of left and right precentral gyrus (PCG) and left and right insula were extracted, as defined in the Automated Anatomical Labeling (AAL) atlas [17]. For further details see Pfurtscheller et al. [5].

### HRV analysis

ECG and respiration were recorded inside the scanner with a sampling rate of 400 Hz. After heart beat detection using fMRI plug-in for EEGLAB [18] and calculation of beat-to-beat interval (RRI) time course (sample rate 4 Hz) the time course was saved as text file and imported to the KUBIOS HRV Premium Package (Kubios Ltd. Finland; version 3.0.2) [19]. Spectral analyses using a Fast Fourier transform algorithm with a window width of 125 s window with 50 % overlap was applied to determine the frequency domain estimates of low frequency (LF) HRV in the two bands 0.06–0.1 Hz (LFa) and 0.1–0.14 Hz (LFb). For statistical analysis, natural log transformed power (log power) and relative power (% power) were used.

### BOLD data processing

For processing of BOLD signals the “Cross-wavelet and Wavelet Coherence” toolbox [20] was used. After bandpass filtering between 0.07 and 0.13 Hz the phase-locking value (PLV) was computed for the two pairs of BOLD signals from PCG and insula and the parameters time delay (TD) and significant length of phase coupling (*%sigbins*) were extracted. For further details see Pfurtscheller et al. [5].

## Results

### Anxiety state and drop rate

State anxiety varied across participants and resting states R1 and R2 in a broad range between 11 (low anxiety) and 29 (rather high anxiety) and significantly declined from R1 (AS1=19.96 ± 4.5) to R2 (AS2= 16.83 ± 4.41) (t(22)=2.39 (p = .026)) (Fig. 1A). Of note, initial state anxiety (AS1) was significantly higher than in the normative sample [13] namely M = 19.96, SD = 4.5 vs. M_normative sample_ = 16.3, t(22) = 3.88, p = 0.001, thus suggesting comparably high levels of anxiety in the first resting state R1.

**Fig. 1:**
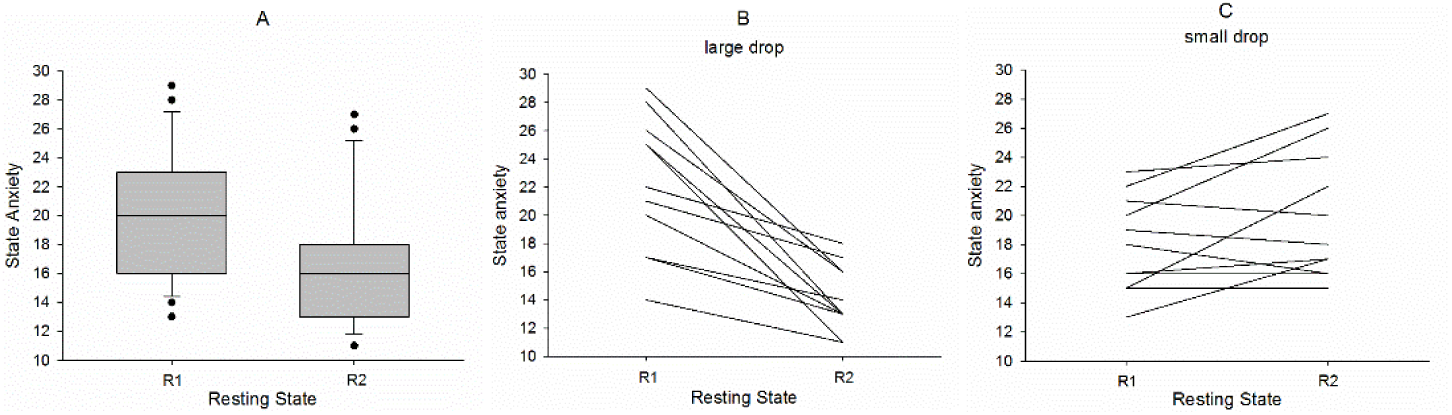
Boxplot depicting the distribution of state anxiety across the resting states R1 and R2 for all 23 subjects (A). Note: whiskers indicate the 10th and 90th percentile. Examples of strong anxiety decline (large drop) in a group of 12 subjects (B) and weak anxiety decline or anxiety increase (small drop) in a group of 11 subjects (C).

An interesting aspect of anxiety processing in consecutive resting states is the drop rate (d). The drop rate is defined as difference of anxiety scores (possible range of AS scores: 10–40) in two selected resting states. For instance d12 defines the drop between resting states R1and R2 (e.g., subject 18: AS1=28 in R1 and AS2= 13 in R2, d12=-15). We found subjects with equal anxiety in R1 and R2 (d12= 0), positive drop rate up to d12 =7 and negative drop rate up to d12=-15. To emphasize the difference between large and small (inclusive zero) drop rate, two diagrams with the corresponding anxiety changes between R1 and R2 are displayed in Fig.1B (large drop) and Fig.1C (small drop).

### Lateralization of phase coupling between BOLD oscillations at 0.1 Hz in precentral gyrus and insula

From each of the 23 individuals, two pairs of PLV parameters (delay and %sigbins), one from each hemisphere, were extracted for both resting states (R1, R2). The grand averages of the parameters delay and %sigbins (mean ± SD), separated for each hemisphere and resting state are summarized in Table 1. In addition, the significance (t-test) of hemispheric differences is indicated. Remarkably, the time delays revealed as significant lateralization with larger nTD values in the left hemisphere most pronounced in R1. The mean duration of significant phase coupling (%sigbins) between BOLD oscillations varied between 38% – 53% and was also larger on the left side. It is noteworthy that the most significant lateralization in R1 was associated with the highest state anxiety in R1.

**Table 1:**
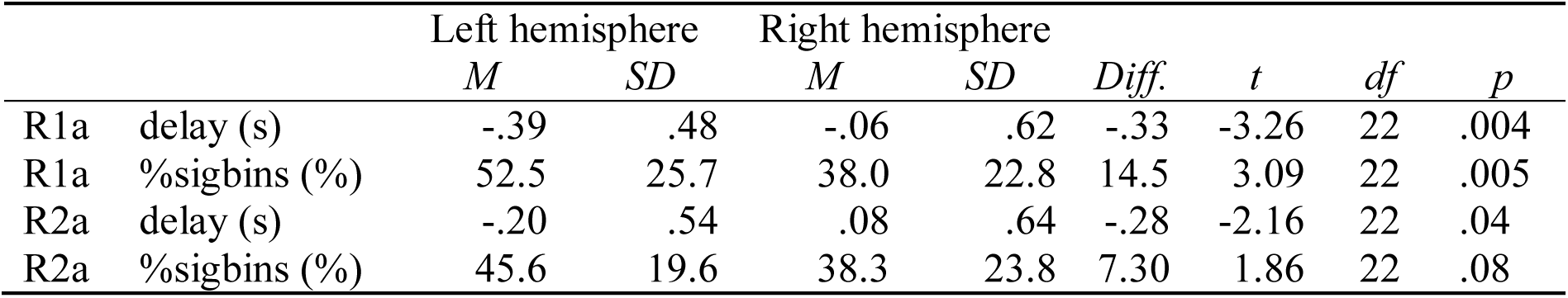
Mean (M), standard deviation (SD) and inter-hemispheric difference (Diff.) of delay and %sigbins in 23 subjects.

|  |  | Left hemisphere |  | Right hemisphere |  | Diff. | t | df | p |
| --- | --- | --- | --- | --- | --- | --- | --- | --- | --- |
|  |  | M | SD | M | SD |  |  |  |  |
| R1a | delay (s) | -.39 | .48 | -.06 | .62 | -.33 | -3.26 | 22 | .004 |
| R1a | %sigbins (%) | 52.5 | 25.7 | 38.0 | 22.8 | 14.5 | 3.09 | 22 | .005 |
| R2a | delay (s) | -.20 | .54 | .08 | .64 | -.28 | -2.16 | 22 | .04 |
| R2a | %sigbins (%) | 45.6 | 19.6 | 38.3 | 23.8 | 7.30 | 1.86 | 22 | .08 |

### Relationship between drop rate and phase coupling of slow BOLD oscillations

For both hemispheres the correlation between drop rate and phase coupling (time delay) of BOLD oscillations in PCG and insula was calculated. This correlation was significant for the left hemisphere (r=0.61, p=0.002) and non-significant for the right side (r=0.33, p=0.129). The diagram with significant correlation between anxiety drop rate and phase coupling of BOLD oscillations (Fig. 2) indicates that different types of anxiety processing can be discriminated. The application of a cluster analysis revealed no clear threshold for the discrimination between two groups, because of their non-homogeneity. Thus, we introduced an arbitrary threshold of d12= -3. The group *large drop* (d12<= -3) contains subjects of high anxiety in R1 and low anxiety in R2 concentrated in the left lower part of Fig. 2 and the group *small drop* (d12>-3) includes subjects displaying nearly constant or even increasing anxiety (right side of Fig. 2). It is important to note, that a nearly stable drop rate was found in subjects with low but also with high anxiety state in R1. The main difference between both subgroups is that the large drop group is associated with only vascular BOLD oscillations (nTD) in the left hemisphere, while the small drop group is associated with a mix of both, vascular (nTD) and neural BOLD oscillations (pTD).

**Fig. 2:**
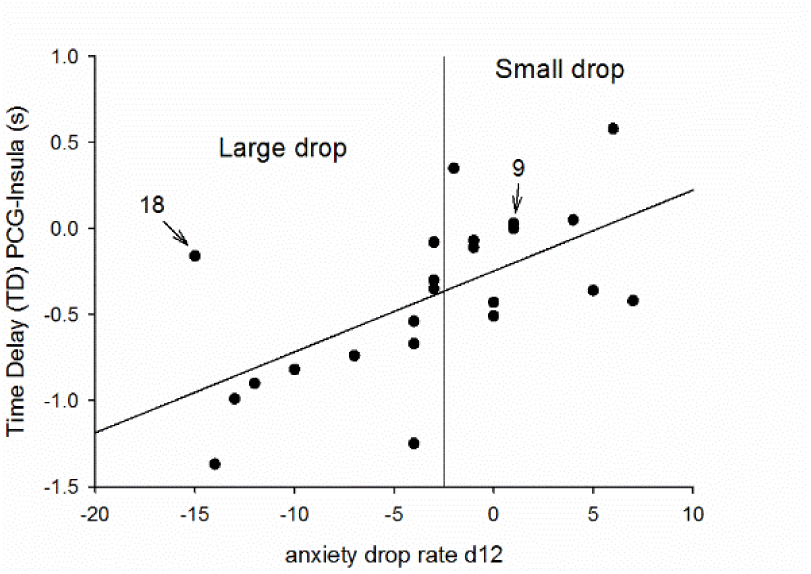
Relationship between drop rate (d12) and significant negative or positive time delay (pTD, nTD) of BOLD oscillations in left precentral gyrus and left insula. Data from resting state R1. The arbitrary threshold is indicated by a horizontal line. Two subjects are indicated by arrows.

To illustrate the trajectories of BOLD, breathing and RRI signals in one member of the small drop group (subject 9R1; indicated by an arrow in Fig. 2), raw signals, spectra and averaged waves are summarized in Fig.3. Because of the lack of triggers in resting state data, RRI peak-triggered averaging was used [7, 21]. To obtain triggers, dominant peaks in the RRI time course are marked (indicated by vertical lines in Fig. 3) and used as trigger to compute average waves of 12-s length (with 6 s before the trigger) for RRI, breathing and BOLD signals. The example depicts spectral peaks at 0.13 Hz in BOLD and RRI signals and a broad peak at 0.3 Hz in the spectrum of respiration. This broad spectral peak underlines the instability of the breathing signal with 2 or 3 breaths between two consecutive RRI peaks, and the averaged breathing wave refers to a phase coupling between slow RRI (0.13 Hz) and breathing (~ 0.3 Hz) oscillations. This example suggests the existence of neural BOLD signals (RRI precedes BOLD oscillations; [7]) with frequencies f > 0.1 Hz in resting states and documents the traveling of neural BOLD waves downwards (descending BOLD oscillations) from precentral gyrus (ROI 1) to insula (ROI 29) within ~ 1.5–0.9 = 0.6 seconds.

**Fig. 3:**
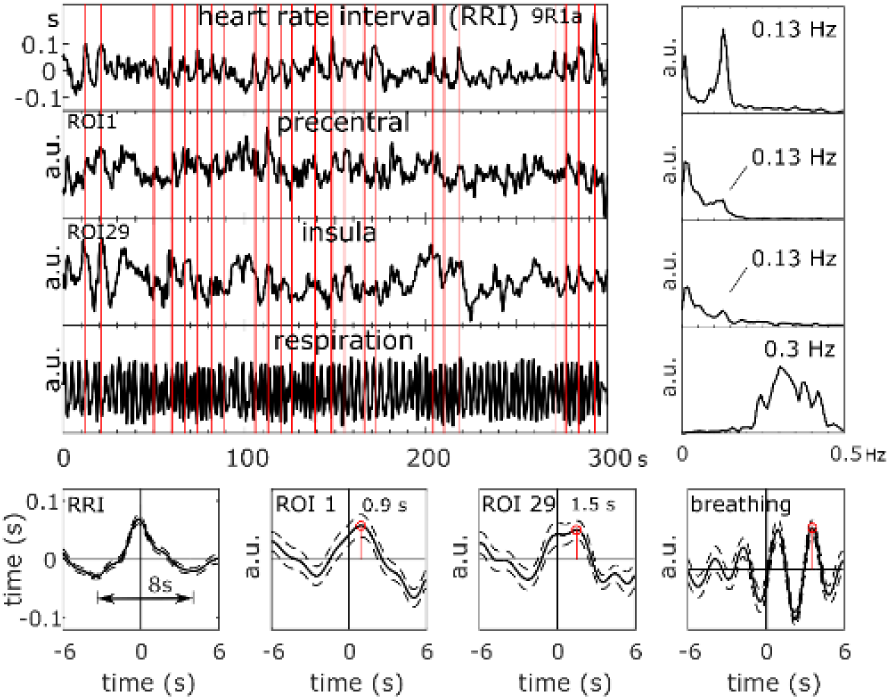
RRI time course, BOLD signals from PCG and insula and respiration (top left), corresponding power spectra (top right), and RRI peak-triggered averages (± SE; bottom) of RRI, BOLD and breathing signals from one subject (9; anxiety score AS1=23) with neural BOLD oscillations (pTD) in resting state R1 and high anxiety. The vertical lines indicate trigger and RRI peak maxima, respectively.

### Relationship between drop rate and HRV in low frequency bands below and above 0.1 Hz

The RRI data of one subject were missing in R2 and were disturbed in R1 in another subject, both belonging to the small drop group. Therefore, only 9 subjects were left for analyses. The results of low frequency (LF) HRV changes from R1 to R2 are summarized in Figs. 4A for all subjects, for subjects with *large drop* (d12<=-3) in Fig 4B and for subjects with *small drop* (d12 >-3) in Fig. 4C separately for LFa (0.06–0.1 Hz) and LFb (0.1–0.14 Hz). The interaction between resting state x component (F(1,19) =8.56, p = 0.009) was significant. This result indicates that both LFa power and LFb power increased from R1 to R2 (Fig.4A), while anxiety significantly decreased (Fig.1A). No significant interaction was found between component (LFa, LFb) x resting state (R1, R2) x group (large and small drop rate) (F1,19)=2.55, p=0.127). However, the interaction between resting state x group was significant (F(1,19) =4.41, p= 0.049) and the interaction between component x group approached significance (F(1,19) = 3.70, p= 0.070). Interestingly, there were between-group (large and small drop) differences in LFa and LFb power from R1 to R2. Whilst LFa power increased significantly (t(11) =2.41, p=0.035) in the large drop group, both components, LFa power (t(8) = 2.69, p= 0.027) and LFb power (t(8) = 3.16, p= 0.013) increased significantly in the small drop group.

**Fig.4:**
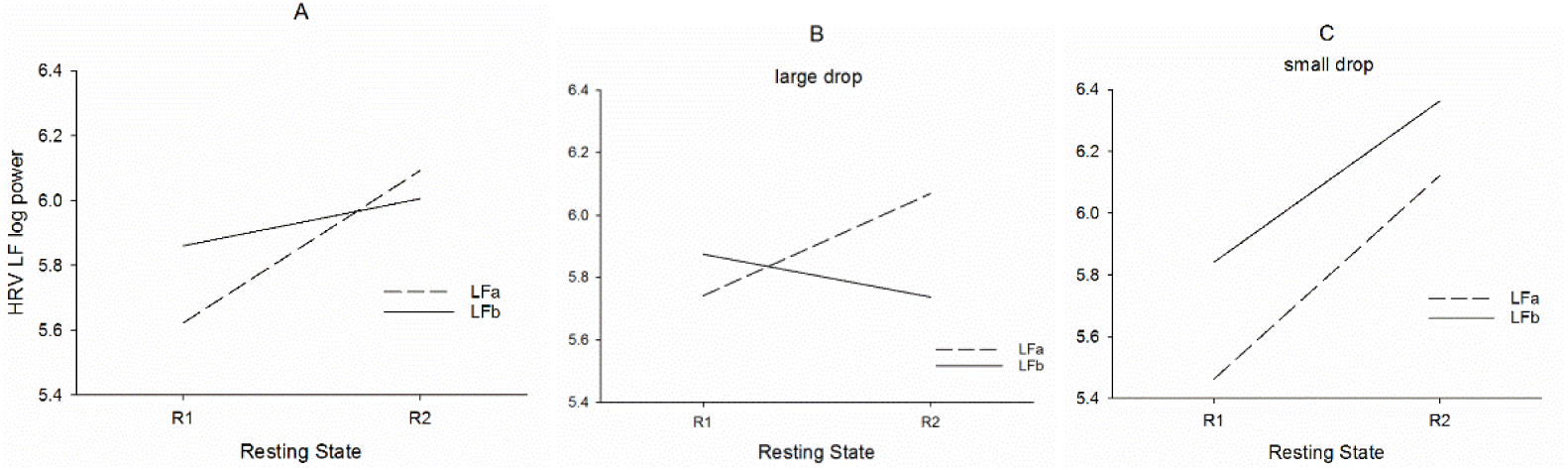
Interactions between resting states (R1, R2) HRV LFa log power (stippled lines) and LFb log power (full lines) for all subjects (A), twelve subjects with *large drop* (B) and nine subjects with *small drop* (C).

### Relationships between neural BOLD oscillations and HRV in two narrow frequency bands below and above 0.1 Hz

In contrast to the left hemisphere, no correlation between drop rate and phase coupling of slow BOLD oscillations was found for the right side, because of the non-dominance of vascular BOLD oscillations and the larger number of subjects with neural BOLD oscillations, respectively. If neural BOLD oscillations are associated with slow neural activity fluctuations, then a positive correlation between neural BOLD oscillations (%sigbins) and HRV LF %power is expected. Such a significant correlation was found, however, only for the LFb band (Fig.4) and emphasizes therewith the existence of central commands preferably above 0.1 Hz.

## Discussion

The discrimination between neural and vascular BOLD oscillations based on PLV calculation depends on several factors. The PLV can either be calculated between BOLD signals and HR interval (RRI) time courses [6,7] or between two BOLD signals [5], whereby the ROI selected for BOLD signal recording is a crucial issue. The latter method is of interest, because no ECG recording is necessary within the scanner and by choosing ROIs in the supply territory of a specific main cerebral artery, the vascular BOLD can be better defined. We monitored the BOLD signal in the supply territories of the left and right middle cerebral artery (MCA), in PCG and insula, two regions of great importance for conscious experience of intending to act [22] and control of cardiac function [23, 24]. The MCA is commonly used to measure cerebral blood flow velocity (CBFv) during rest and complex cognitive tasks as it supplies 80% of each respective cerebral hemisphere’s frontal lobe [25].

### MRI-related anxiety and drop rate

It is known that anxiety is elevated during the first scan and drops to normal levels later [3]. Surprisingly, the anxiety change (drop rate) between two resting states separated by ~ 30 min shows considerably variation as documented in Fig. 1B and 1C, either in form of a sharp decline (large drop) or even a moderate increase (small drop). These different trajectories in state anxiety are particularly interesting and suggest that different person-specific mechanism of anxiety regulation might have been activated.

### Relationship between anxiety drop rate and phase coupling of slow BOLD oscillations in the territory of the left MCA

Spontaneous BOLD oscillations around 0.1 Hz can be generated in parts by a complex interplay of slow cerebral blood volume and CBFv oscillations [26] and by intrinsic neural activity fluctuations [8]. The former is closely associated to the CBFv measured by transcranial Doppler sonography in the main cerebral arteries and phase-coupled with blood pressure waves measured via infrared finger plethysmography [27, 28]. In healthy subjects and rest, the CBFv of both MCAs oscillates nearly synchronously and demonstrates hemispheric symmetry [25, 28]. Intrinsic neural activity fluctuation around 0.1 Hz can be observed in EEG or ECoG recordings and have been reported in the beta/alpha power at EEG electrodes over sensorimotor areas [21] and in the beta and gamma power at multiple electrode sites placed in human posteromedial cortex [29]. The close relationship between gamma power and BOLD oscillations (neural BOLD) was documented impressively by Mateo et al. [8] via entrainment of arteriole vasomotor fluctuations by neural activity.

Both types of slow BOLD oscillations have a different origin and can thus be composed by different frequency components. Our assumption was that a discrimination should be possible by their direction of spreading, vascular BOLD oscillations are ascending in the supply territory of MCA and neural BOLD oscillations are descending from higher centers in direction to the cardiovascular centers in the brain stem ultimately modulating heart rate [11]. The trend toward larger nTD magnitudes with higher drop rate (Fig. 2) may be interpreted as accumulating dominance of vascular BOLD oscillations. The opposite, the trend toward larger pTD magnitudes may be indicative for a accumulating dominance of neural BOLD oscillations. Both, the significant correlation between drop rate (d12) and phase coupling (TD) in the left hemisphere (Fig. 2) and the significant hemispheric asymmetry with the predominance of nTD in the left side (Table 1) provide strong hints toward a cerebral blood flow increase in the territory of the left MCA, which could constitute a precondition for a successful processing of high anxiety states.

Anxiety and/or fear is not only accompanied by activation of the left amygdala [30, 31] but also of the left insular cortex [4]. Dennis et al. [4] reported a 6-min resting state study on young and adult participants with scanning followed by questionnaires to assess their mood and thoughts during the scan. Both groups showed increased connectivity between left insular cortex and the default mode network [32] with increased anxiety. This left-sided insular activation during task-free periods suggests that increased state anxiety is reflected in resting-state functional connectivity. Positron emission tomography (PET) revealed an increase of cerebral blood flow during emotion induction in left orbitofrontal and left anterior insular cortices, which was correlated with HF-HRV [33]. Common in these reports [30,31,32] is the involvement of structures in the left hemisphere preferentially involved in regulating vagal tone. Together these findings suggests an increase in cerebral metabolic activity – but also its by-product carbon dioxide – in left amygdala, left insula and related structures especially during negative emotion processing. This in turn leads to a local increase of CBF to supply the areas with oxygenated blood and remove waste products [34]. The lateralized increase of cerebral blood flow circulation in the left MCA in the majority of subjects with high state anxiety is therefore not unexpected and contributes to a concomitant increase in low frequency HRV.

### Relationship between anxiety drop rate and HRV in low frequency bands below and above 0.1 Hz

An interesting finding refers to the relationship between state anxiety and HRV. A combined BOLD-HRV study in patients with posttraumatic stress disorders (PTSD) and healthy controls revealed higher low frequency (LF) HRV and increased connectivity between left amygdala and periaqueductal gray in controls [35]. In this case the standard LF band (0.04–0.15 Hz; Task Force Guidelines 1996) was used. Therefore, it is assumed that the LF HRV should display an increase from R1 to R2 in healthy scanner-naive subjects.

All studies using HRV as important biomarker for stress, pain, anxiety and other unpleasant emotions [10, 11, 33, 36, 37] report on two major components, a high frequency component (0.15–0.4 Hz) and a low frequency component (0.01–0.15 Hz or 0.04–0.15 Hz). To our knowledge there is no report differentiating between different low frequency components, although the LF component depends on a mixture of both parasympathetic and sympathetic autonomic influences [13]. The existence of two distinguishable low frequency bands is confirmed by a spectral analysis of HR and BP signals [12]. They reported on two principal frequency components at 0.08 Hz and 0.12 Hz, however, their underlying generating mechanisms were not elucidated. The discrimination between HRV frequency bands below (LFa: 0.06–0.1 Hz) and above 0.1 Hz (LFb: 0.1–0.14 Hz) is of multiple interest and leads among others to the following question: Is any one of these frequency bands predominant in the context of the *pacemaker theory* proposed by Julien [38] assuming the existence of rhythmic central commands? Can the discrimination between two LF HRV components help to differentiate between diverging anxiety processing strategies? Is one band more indicative for a parasympathetic influence?

Considering the different and variable patterns associated with a large drop of anxiety in one group and no drop or even an anxiety increase in the other group (examples see Fig. 1B and C) it is not astonishing that no significant interaction was found between resting state, group and component. The significant interaction between resting state and component documents the expected increase of LF HRV in connection with a decline in anxiety from R1 to R2 (Fig. 4A). Noteworthy, there was also a significant interaction between resting state and group (Fig. 4A and C) with a significant increase of both LF components from R1 to R2 in the small drop group and an increase of only LFa power in the large drop group. Based on the HRV analyses this can be interpreted that frequency components between 0.1–0.14 Hz may play a minor role in subjects with large anxiety drop, however a major role in subjects with small drop.

Considering both the HRV analysis and the correlation between drop rate and BOLD phase-coupling, it might be speculated that one neural strategy of anxiety processing is based on enhanced vascular BOLD oscillations (*baroreflex theory*) predominant in the band 0.06–0.1 Hz, and the other strategy is based on reinforced rhythmic central commands (*pacemaker theory*) with a frequency preference in the 0.1–0.14 Hz band. These central commands originating in prefrontal cortex control cardiac activity via vagus nerve activation [24]. Herewith the parasympathetic influence in the 0.1–0.14 Hz band may be explained. Further research is necessary however, to verify or falsify this hypothesis.

### Relationships between neural BOLD oscillations and low frequency band HRV

The majority of neural BOLD oscillations (pTD) was found in the right hemisphere in R1. Such descending neural BOLD oscillations were observed in twelve subjects. If these oscillations are associated with central commands projecting to the cardiovascular nuclei in the brain stem and are responsible for the fast modulation of HR, then there should emerge a correlation between the length of significant phase coupling (%sigbins) in descending BOLD oscillations and LF HRV. The results in Fig. 5 indicate that such a correlation is likely, however, only for the LFb band (0.1–0.14 Hz). This finding not only confirms the reported phase-coupling between neural BOLD oscillations and RRI signals [7], but gives evidence that neural BOLD oscillations are not only restricted to frequency components below 0.1 Hz [39] and shows that central commands operates dominantly at frequency component above 0.1 Hz.

**Fig. 4:**
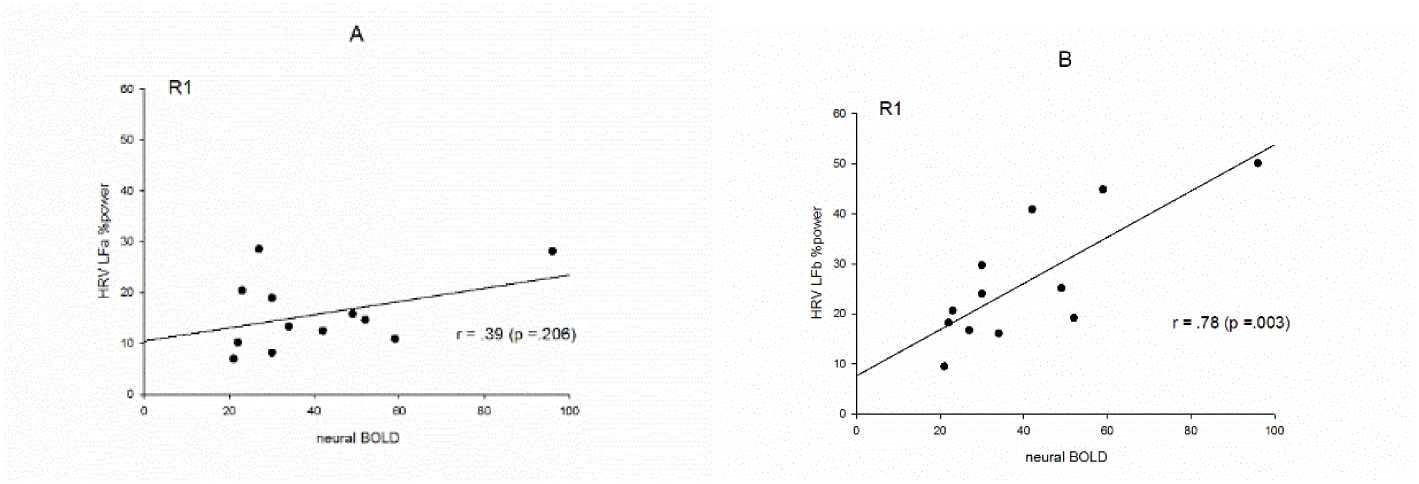
Relationships between HRV LF relative (%) power and percentage of significant phase coupling (%sigbins) between PCG and insula in the right hemisphere for neural BOLD oscillations (pTD). Low frequency %power in the band 0.06–0.1 Hz (LFa%) (A) and low frequency %power in the band 0.1–0.14 Hz (LFb%) (B).

Conclusion:

1. Healthy, scanner-naïve individuals displayed MRI-related state anxiety ether with a strong anxiety decline (large drop) between two resting states at intervals of ~ 30 minutes or a moderate anxiety change (small drop). This suggests that there are different mechanism of anxiety processing within the sample.
2. Through measuring the phase-coupling between intrinsic BOLD oscillations at 0.1 Hz in precentral gyrus and insula it is not only possible to discriminate between vascular and neural BOLD oscillations, but also between subjects with either a large or a small anxiety drop. Subjects with large anxiety drop revealed a hemispheric asymmetry especially during the first resting state with a predominance of vascular BOLD oscillations at 0.1 Hz in the left hemisphere and subjects with a small drop showed a dominance of intrinsic neural BOLD oscillations at 0.1 Hz.
3. The analysis of within-scanner HRV revealed a pronounced increase of low frequency power between both resting states. This power increase was dominant in the band 0.06–0.1 Hz in the large drop group and in the band 0.1–0.14 Hz in the small drop group and suggests that rhythmic central commands modulating heart rate via vagus nerve activation operate dominantly at frequencies above 0.1 Hz.

## Limitations

One problem is the small group size and their non-homogeneity with a relatively great number of subjects with moderate drop rate and small non-reliable pTD or nTD (%sigbins < 20%). Another limitation is the use of only two pairs of BOLD oscillations in the territory of MCA. Nevertheless, it is surprising that both groups could be discriminated based on ascending or descending BOLD oscillations at 0.1 Hz. Improvement is expected, if BOLD signals from supply territories of MCA and anterior cerebral artery (medial prefrontal cortex) are investigated and mean PLVs between BOLD signals are calculated.

An additional point is that patterns of PLV (nTD or pTD) are not subject-specific and not stable across resting states. They can change from nTD in resting state R1 to pTD in R2 and vice versa. This unstable behavior was reported already and denoted by “switching phenomenon” [6]. It could be speculated, that in cases of a high anxiety level not only one, but both strategies (blood flow increase and central commands) are activated. An example for this is for example subject 18R1 classified as member of the large drop group and marked as outlier in Fig. 2. This subject displayed two peaks in the HRV spectrum (at 0.09 Hz and 0.11 Hz) with LFb power > LFa power in R1 and LFb power < LFa power in R2. The dominance of HRV power in the 0.1–0.14 Hz band in the initial resting state R1 with high anxiety (AS=28) suggests that in addition to the blood flow increase also strong central commands were active.

A problem could also be the high inter-and intra-subject variability of breathing patterns and the selection of LF and HF HRV bands. Breathing is regulated either by respiratory neurons in medulla oblongata and pons (metabolic breathing) or from higher centers in the cerebral cortex (behavioral breathing) and can be conscious (thinking about the breath) or unconscious [40, 41]. A special type of respiration is breathing at 6/min (0.1 Hz) which approximates the resonance frequency of the human baroreflex loop [8, 42]. In our study no instruction was given about the type of respiration. The majority of participants displayed spontaneous breathing at a rate of 0.2–0.35 Hz, a minority displayed breathing at 0.1 Hz (6/min) especially in the first resting state R1 where the state anxiety was highest. While breathing at a rate between 0.2–0.35 Hz belongs to the HF HRV band, resonance breathing at ~ 6/min contributes to the LF band. Due to this variability of the respiration it is not astonishing that the interaction of HRV LFa and LFb power between resting state (R1, R2) and group (large drop, small drop) reached just significance (p < 0.049). One solution of the problem could be a controlled breathing at a subject-specific rate in the HF band.

## Acknowledgments

We like to thank Thomas Zussner and Karl Koschutnig, University of Graz, for support in data acquisition.

<u>Author Contributions:</u>

Conceptualization: GP, AS

Data curation: CB, DF

Formal analysis: AS, CB, AA

Investigation: AA Methodology: GP, AA

Project administration: GP

Resources: GP, AS

Software: AA, CB, CA, JB

Supervision: GP, AA, AS

Validation: GP, AA

Visualization: AA, AS, CA

Writing-original draft: GP

Writing-review & editing: GP, AA, AS

## References

1. Katz RC, Wilson L, Frazer N. Anxiety and its determinants in patients undergoing magnetic resonance imaging. J Behav Ther Exp Psychiatry. 1994; 25(2):131–4.

2. Dewey M, Schink T, Dewey CF. Claustrophobia during magnetic resonance imaging: cohort study in over 55.000 patients. J Mag Res Imag. 2007; 26: 1322–1327.

3. Chapman HA, Bernier D, Rusak B. MRI-related anxiety levels change within and between repeated scanning sessions. Psych Res: Neuroimaging. 2010; 182:160–164.

4. Dennis EL, Gotlib IH, Thompson PM, Thomason ME. Anxiety modulates insula recruitment in resting-state functional magnetic resonance imaging in youth and adults. Brain Connectivity. 2011; 1(3): 245–254.

5. Pfurtscheller G, Schwerdtfeger A, Brunner C, Aigner C, Fink D, Brito J, et al. Distinction between neural and vascular BOLD oscillations and intertwined heart rate oscillations at 0.1 Hz in the resting sate and during movement. PloSOne. 2017 Jan4; e0168097

6. Pfurtscheller G, Schwerdtfeger A, Seither-Preisler A, Brunner C, Aigner CS, Calito J, et al. Brain-heart communicatio: Evidence for “central pacemaker” oscillations with a dominant frequency at 0.1 Hz in the cingulum. Clin Neurophys. 2017; 128:183–193.

7. Pfurtscheller G, Schwerdtfeger A, Seither-Preisler A, Brunner C, Aigner CS, Calito J, et al. Synchronisation of intrinsic 0.1-Hz blood-oxyden-level-dependent (BOLD) oscillations in amygdala and prefrontal cortex in subjects with increased state anxiety. Eur J Neurosci. 2018; doi: 10.1111/ejn.13845

8. Mateo C, Knutsen PM, Tsai PS, Shih AY, Kleinfeld D. Entrainment of arteriole vasomotor fluctuations by neural activity is a basis of blood-oxygenation-level-dependent “resting-state” connectivity. Neuron. 2017; 96:1–13.

9. Mather M, Thayer J. How heart rate variability affects emotion regulating brain networks. Current Opinion Behavioral Science. 2018; 19:98–104.

10. Appelhans BM, Luecken LJ. Heart rate variability and pain: Associations of two interrelated homeostatic processes. Biol Psychology 2008; 77:174–182

11. Thayer JE, Ahs F, Fredrikson M, Sollers JJ, Wager TD. A meta-analysis of heart rate variability and neuroimaging studies: implication for heart rate variability as a marker of stress and health. Neurosc Behav Rev. 2012; 36:747–756.

12. Kuusela TA, Kaila, TJ, Kähönen M. Fine structure of the low-frequency spectra of heart rate and blood pressure. 2003; BMC Physiol. 13: 3–11.

13. Laux L, Hock M, Bergner-Koether R, Hodapp V, Renner KH, Merzbacher G. Das State-Trait-Angst-Depressions-Inventar [The State-Trait Anxiety-Depression Inventory]. Goettingen Hogrefe; 2013.

14 Spielberger CD, Gorssuch RL, Lushene PR, Vagg PR, Jacobs G. Manual for the State-Trait Anxiety Inventory. Palo Alto CA Consulting Psychologists Press Inc; 2009.

15. Moeller S, Yacoub E, Olman CA, Auerbach E, Strupp J, Harel N, et al. Multiband multislice GE-EPI at 7 Tesla, with 16-fold acceleration using partial parallel imaging with application to high spatial and temporal whole-brain fMRI. Magn Reson Med. 2010;63:1144–53.)

16. Chao-Gan Y, Yu-Feng Z. DPARSF: a MATLAB toolbox for “pipeline” data analysis of resting-state fMRI. Front Syst Neurosci. 2010; 4(May):13.

17. Tzourio-Mazoyer N, Landeau B, Papathanassiou D, Crivello F, Etard O, Delcroix N, et al. Automated Anatomical Labeling of Activations in SPM Using a Macroscopic Anatomical Parcellation of the MNI MRI Single-Subject Brain. Neuroimage. 2002;15(1):273–89.

18. Niazy RK, Beckmann CF, Iannetti GD, Brady JM, Smith SM. Removal of fMRI environment artifacts from EEG data using optimal basis sets. Neuroimage. 2005; 28(3):720–37.

19. Tarvainen MP, Niskanen J-P, Lipponen JA, Ranta-aho PO, Karjalainen PA. Kubios HRV – Heart rate variability analysis software. Comput Methods Programs Biomed. 2014;113(1):210–20.

20. Grinsted A, Moore JC, Jevrejeva S. Application of the cross wavelet transform and wavelet coherence to geophysical time series. Nonlinear Process Geophys. 2004;11(5/6):561–6.

21. Pfurtscheller G, Daly I, Bauernfeind G, Müller-Putz GR. Coupling between intrinsic prefrontal HbO2 and central EEG beta power oscillations in the resting brain. PLoS One. 2012 Jan;7(8):e43640.

22. Haggard P. Conscious intention and motor cognition. Trends Cog Sciences. 2005; 9(6): 290–295.

23. Verberne AJ, Owens NC. Cortical Modulation of the Cardiovascular System. Prog Neurobiol. 1998; 54(2):149–68.

24. Thayer JF, Lane RD. Claude Bernard and the heart-brain connection: Further elaboration of a model of neurovisceral integration. Neurosci Biobehav Rev. 2009;33(2):81–8.

25. Harwood AE, Greenwood M, Shaw TH. Transcranial doppler sonography reveals reduction in hemispheric asymmetry in healthy older adults during vigilance. Front Aging Neurosci. 2017; Feb 2017, Vol 9, Artice 21

26. Obrig H, Neufang M, Wenzel R, Kohl M, Steinbrink J, Einhäupl K, et al. Spontaneous low frequency oscillations of cerebral hemodynamics and metabolism in human adults. Neuroimage. 2000; 12(6):623–39.

27. Aaslid R. Noninvasive transcranial Doppler ultrasound recording of flow velocity in basal cerebral arteries. J Neurosurg. 1982; 57(6): 769–774.

28. Diehl RR, Linden D, Lucke D, Berlit P. Phase Relationship Between Cerebral Blood Flow Velocity and Blood Pressure : A Clinical Test of Autoregulation. Stroke. 1995; 26(10):1801–4.

29. Foster BL, Parvizi J. Resting oscillations and cross-frequency coupling in the human posteromedial cortex. Neuroimage. 2012 Mar;60(1):384–91.

30. Phelps EA, O’Connor KJ, Gatenby JC, Gore JC, Grillon C, Davis M. Activation of the left amygdala to a cognitive representation of fear. Nature Neurosc. 2001; 4:437–441.

31. Baas D, Aleman A, Kahn RS. Lateralization of amygdala activation: a systematic review of functional imaging studies. Brain Res Rev. 2004; 45:96–103

32. Raichle ME, MacLeod AM, Snyder AZ, Powers WJ, Gusnard DA, Shulman GL. A default mode of brain function. PNAS. 2001;98: 676–682.

33. Lane RD, McRae K, Reiman EM, Chen K, Ahern GL, Thayer JF. Neural correlates of heart rate variability during emotion. Neuroimage. 2009; 44:213–222.

34. Aaslid R. Transcraneal Doppler examination techniques. In: Aaslid R, editor. Transcraneal Doppler Sonography. Springer Berlin 1986. pp 35–59.

35. Thome J, Densmore M, Frewen PA, McKinnon MC, Theberge J, Nicholson AA, et al. Desynchronisation of autonomic response and central autonomic network connectivity I posttraumatic stress disorder. Human Brain Mapping 2017; 38(1): 27–40.

36. Lagos L, Vaschillo E, Vaschillo B, Lehrer P, Bates M, Pandina R. Heart rate variability biofeedback as a strategy for dealing with competitive anxiety: a case study. Biofeedback. 2008; 36:109–115.

37. Williams DP, Feeling NR, Hill LK, Spangler DP, Koenig J, Thayer JF. Resting heart variability, facets of rumination and trait anxiety: implications for the perseverative cognition hypothesis. Front Hum Neurosci. 2017; 31 Oct

38. Julien C. The enigma of Mayer waves: Facts and models. Cardiovasc Res. 2006; 70:12–21.

39. Snyder AZ, Raichle ME. A brief history of the resting state: the Washington University perspective. Neuroimage. 2012; 62, 902–910.

40. Homma I, Massaoka Y. Breathing rhythms and emotions. Exp Physiology. 2008; 93.9: 1011–1021.

41. Brewer JA, Garrison KA. The posterior cingulate cortex as a plausible mechanistic target of meditation: findings from neuroimaging. Ann N Y Acad Sci. 2013; 1–9

42. Lehrer P. How does heart rate variability biofeedback work? Resonance, the baroreflex, and other mechanisms. Biofeedback. 2013; 41:26–31.

